# Horizontal gene transfer-mediated bacterial strain variation affects host fitness

**DOI:** 10.1101/2020.09.24.311167

**Authors:** Yun Wang, Franz Baumdicker, Sven Kuenzel, Fabian Staubach

**Author notes:** **Corresponding author:** Fabian Staubach.

## Abstract

How microbes affect host fitness and environmental adaptation has become a fundamental research question in evolutionary biology. We tested for associations of bacterial genomic variation and *Drosophila melanogaster* offspring number in a microbial Genome Wide Association Study (GWAS). Leveraging strain variation in the genus *Gluconobacter*, a genus of bacteria that are commonly associated with *Drosophila* under natural conditions, we pinpoint the thiamine biosynthesis pathway (TBP) as contributing to differences in fitness conferred to the fly host. By tracing the evolutionary history of TBP genes in *Gluconobacter*, we find that TBP genes were most likely lost and reacquired by horizontal gene transfer (HGT). We suggest that HGT might contribute to microbiome flexibility and speculate that it can also more generally contribute to host adaptation.

## Introduction

Microbes are important drivers of host phenotype and evolution [1]. Benefits derived from microorganisms can facilitate the occupation of new ecological niches [2–5] and microbial effects on host phenotypes and fitness can spur adaptive processes [6–14]. Changes in the effects of microbes on host fitness can alter interactions along the parasitism mutualism continuum [6, 15–18], thus affecting the evolutionay trajectories of the partners. The importance of microbes in evolution and health of higher organisms has sparked a search for the molecular underpinnings of how microbes affect host phenotype.

In this search, microbial Genome Wide Association Studies (GWAS) are an important tool [19–23]. The principle behind a microbial GWAS is to establish a link between traits and genetic variation of microbes by the means of GWAS. By testing for association between host traits and microbial genomic variation, Chaston et al. [24] introduced a particularly helpful approach to unravel how microbes affect host phenotypes [22, 24]. The authors measured host phenotypes, here from *Drosophila melanogaster* that were mono-associated with several microbial isolates. Differences in host phenotype were then associated with the presence and absence of genes in the microbial isolates. By applying this approach, it was found that genes that play a role in glucose oxidation in bacteria affect *D. melanogaster* triglyceride levels [24] and that bacterial methionine and B vitamins are important for starvation resistance [25] as well as life span [26].

For targetting host phenotypes with microbial GWAS, model systems that allow the generation of axenic hosts that can successively be associated with individual microbial isolates are particularly useful [24]. One such model system is *D. melanogaster* and its bacterial microbiome. Techniques for the generation of gnotobiotic flies are readily available and standardized measurements of phenotypes exist. Affected phenotypes include the life history traits development time, fecundity, and life span as well as size of the adults [14, 26–33]. These traits are directly related to fitness, emphasizing the potential importance of microbes in host evolution and adaptation. Microbes often affect fitness related traits by provisioning nutrients. These nutrients include vitamins, amino acids, lipids, and trace elements [24, 34–39]. Nutrient provisioning is a recurring theme in macrobe–microbe interactions that are adaptive for the host [40–42].

The acquisition of nutrients from microbes need not rely on microbes that live inside the host. Instead, nutrients can also be acquired by harvesting or preying upon microbes that live outside the fly and subsequent digestion [35, 43–45]. Furthermore, bacteria have been identified that affect *D. melanogaster* phenotypes by increasing the ability for nutrient uptake [46] or metabolizing components of the food substrate, and thus modulating its nutrient content [24]. Interestingly, the metabolic potential to produce nutrients that affect fly fitness differ between closely related microbes and so do the effects on fly phenotype and fitness [26, 29, 32, 47–51]. These findings contribute to the notion that microbial variation at taxonomically low levels is not only important for human [15], mouse [52], and plant [18] hosts, but also for *Drosophila* [53].

Because variation between closely related bacteria is important for the interaction of the host and its microbiome, it is also important to consider closely related microbes in studies that aim at elucidating the molecular underpinnings of host-microbe interaction. At the same time, using the pan-genome of closely related microbes in GWAS might offer particular power to the approach: on a similar genomic background, microbial genes that affect the host are easier to identify. For studies that are aimed at better understanding host-microbe interaction in an evolutionary context, it is also important to consider microbes that are associated with the host under natural conditions and if possible, to measure evolutionary relevant host phenotypes in a natural or near natural environment. Finally, tracing the evolutionary changes of the genomic elements that affect host fitness can help us to gain deeper insights into how host-microbe interaction evolves.

We aimed our study at better understanding whether and how fly fitness is affected by its natural microbiome by a microbial GWAS. In order to increase the power of the approach and consider variation at low taxonomic levels, we concentrated on variation within a taxonomically restricted group of bacteria. Therefore, we focused our study on *Gluconobacter*, a bacterial genus that is commonly associated with *D. melanogaster* under natural conditions [54–57]. We assessed fitness parameters on grape juice based fly food as a near natural food source. In order to better understand how microbial effects on host fitness evolve, we traced the evolutionary events that lead to changes in bacteria-mediated host fitness.

## Materials and Methods

### Fitness assays

Canton-S stocks were kept at 25 °C on food prepared following the Bloomington *Drosophila* Stock Center ‘Cornmeal Molasses and Yeast Medium’ (532 ml water, 40 ml molasses, 6.6 g yeast. 32.6 g cornmeal, 3.2 g agar, 2.2 ml propionic acid and 7.6 ml Tegosept). To generate axenic flies, embryos were collected and washed in PBS, dechorionated in 50% bleach for 2-3 mins, and rinsed in sterile PBS for 1 min. Embryos were placed in sterilized food bottles under a sterile workbench and maintained at 25 °C under a 12:12 light cycle in axenic condition for 3 weeks during which the flies had time to hatch and mate. One axenic female from these bottles was used per vial in the fitness assay. For the fitness assay, bacterial cultures were grown in liquid YPD medium for 48-72 hours and normalized to the same optical density (OD600 = 0.6). 150 μl of OD normalized medium were added directly on 10ml sterile grape juice food (667 ml water, 333 ml Jacoby white grape juice, 8 g yeast, 50 g cornmeal, 10 g agar, 3 ml propionic acid). Axenic females were transferred to the vial immediately afterwards. We prepared two control treatments. First, we added sterile YPD medium to the food as axenic control. Second, we used conventionally reared flies homogenized in YPD as inoculum. We tracked the number of pupae per vial for 16 consecutive days after infection. On the last day, flies were counted, collected and weighed. All offspring were weighed together in one Eppendorf tube for each replicate and weight per fly was calculated. All fitness related measurements were done blind. That means the vials were given random numbers and only after the measurements were taken, the bacterial strain ID was connected to the result. For the thiamine supplement experiment, we added 1 μg/ml thiamine to the food described above. That concentration has proven effective for phenotypic rescue in [37]. All statistical analyses were performed in R and can be found in supplementary script S1.

### Bacterial loads and contamination control

Fly offspring from the fitness assays were stored in PBS/glycerol mixture at −80 °C for later contamination control and the counting of colony forming units (CFUs). 3-6 replicates per bacterial isolate were picked for CFU counting. For CFU counting, samples of 3 to 5 offspring were homogenized with a pestle in 300 μl of PBS. The homogenates were plated on YPD agar medium. Plates were incubated for 48 hours. CFU counts were done visually or with the OpenCFU software [58], Table S4). Plates for CFU counting were also inspected for colony morphology and colony color that could indicate potential contamination, with negative results. All homogenates were plated on antibiotic YPD agar medium (with 100 μg/ml kanamycin or ampicillin) for assessing yeast contamination. No yeast colonies were observed except in the control replicates in which conventional lab microbiota were used. To further assess potential bacterial contaminants during our experiment, we quantified the relative abundance of target isolates on fly offspring using 16S rRNA gene sequencing. In short, DNA was extracted from pools of 3-5 offspring for 3-6 replicates per bacterial isolate after the experiment, including the replicates with the highest and lowest offspring number. The V4 regions of the bacterial 16S rRNA gene were amplified and sequenced on an illumina MiSeq sequencer following [56, 59]. Sequencing data were analyzed using mothur [60] (See supplementary script S2 for all commands executed). The relative abundance of target 16S rRNA gene sequences for mono-associated isolates was calculated. The average relative abundance of target 16S sequences was over 88% (Figure S8A) in the initial experiment. Only in (6 out of 66) replicates the relative abundance was below 75%, including 3 cases of *P. sneebia* that showed very low bacterial loads. For the thiamine treatment the target bacteria were significantly enriched in the microbial community (Figure S8B).

### Bacterial isolates, genome sequencing and assembly

We sequenced, assembled, and annotated draft genomes of eleven bacteria and added genome data for six bacteria from public databases (Table S1). Nine strains were isolated from wild-caught *Drosophila* collected in the San Francisco Bay Area (California, USA). Isolates were cultured in YPD for standard phenol-chloroform DNA extraction. Bacterial genomes were sequenced using Illumina MiSeq technology and assembled with the A5 MiSeq assembler [61]. Completeness and contamination were assessed with checkM v1.1.2 [62], using standard settings. Assembly statistics were generated with QUAST v5.0.2 [63]. Annotation was performed with prokka v1.1 [64] or imported from GenBank. Average nucleotide identity (ANI) was computed with fastANI (v0.1.2). New isolates were taxonomically classified, using GTDBtk (v0.1.4) [65]. FastANI and GTDBtk were run on the kbase web interface [66].

### Pan-genome clustering and phylogenetic trees

Genomes were analyzed using the panX analysis pipeline [60] with standard parameters (script S3). Genes were grouped into 11269 clusters of homologous sequences, including clusters with a single gene. Thereby the presence and absence of each gene cluster in the 17 genomes was estimated. Based on the alignments of all single-copy genes that are present in all 17 genomes, panX reconstructs a phylogenetic tree. For the phylogeny, FastTree 2 [67] and RaxML [68] are applied to all variable positions from these alignments. For the phylogeny of thiE, nucleotide sequences were aligned using MUSCLE v3.8.425 [99] and the tree was built using MrBayes 3.2.6 [70] as incorporated in the Geneious software suit v1.1 (Biomatters ltd.).

### Microbial pan-genome-wide association study

We calculated the gene presence absence association score (PA score) between each predicted cluster of homologous genes and fly offspring number. I.e. if D_g_ is the difference between the mean fly offspring for strains with and without gene g, σ is the standard deviation of fly offspring and n_g_ is the number gene gains and losses as inferred from the phylogeny. The association score is given by 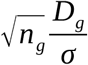. Three alternative association scores from treeWAS [21] and the corresponding model based p-values were calculated. Association scores based on the presence and absence of genes are prone to false positives because genome wide linkage results in strongly correlated presence and absence of genes. Panx and treeWAS reduce this effect by taking the reconstructed ancestral gene gain and loss events into account.

## Results

We performed a microbial GWAS for the number of offspring produced by females that were mono-associated with 17 bacterial isolates from genera that co-occur with *Drosophila melanogaster* in its natural environment. *Gluconobacter* was represented by 13 isolates. Two additional isolates were from the genus *Acetobacter*. Species from this genus can benefit *Drosophila* development [28]. One isolate was *Commensalibacter intestini* that might have a probiotic function in *D. melanogaster* [71] and is enriched in flies over substrate in wild-caught flies [57]. As an outgroup and to get a baseline for the fitness effect of an ingested pathogen, we added *Providencia sneebia*, that is highly pathogenic when entering the hemolyph [72]. All bacterial genomes analyzed were >99% complete with the exception of *P. sneebia* (>96%, Table S1). The mean number of offspring varied significantly between flies mono-associated with different isolates (*P* = 4.2 x 10^-15^, Kruskal-Wallis-Test, Figure 1A) up to a 2.8-fold difference between *Gluconobacter morbifer* and *Gluconobacter sp.* P5H9d. Differences between bacterial strains also explained a significant proportion of variation in offspring number when we accounted for bacterial loads per fly (*P* = 1.4 x 10^-4^, linear model), suggesting that not only bacterial biomass affects fly fitness. Presence-absence patterns of 11269 genes were tested for association with the number of offspring that mono-associated females produced using the PA method [73]. Associations were confirmed by permutation tests and TreeWAS [21] (Figure S1, Table 1, Table S2). The six highest PA scores depended strongly on presence-absence patterns between the closely related strains P1C6b, DSM2003, DSM2343, and DSM3504 (mean ANI = 95.5%) in the branch that includes *G. morbifer* (Figure S1 and accompanying text).

**Figure 1.**
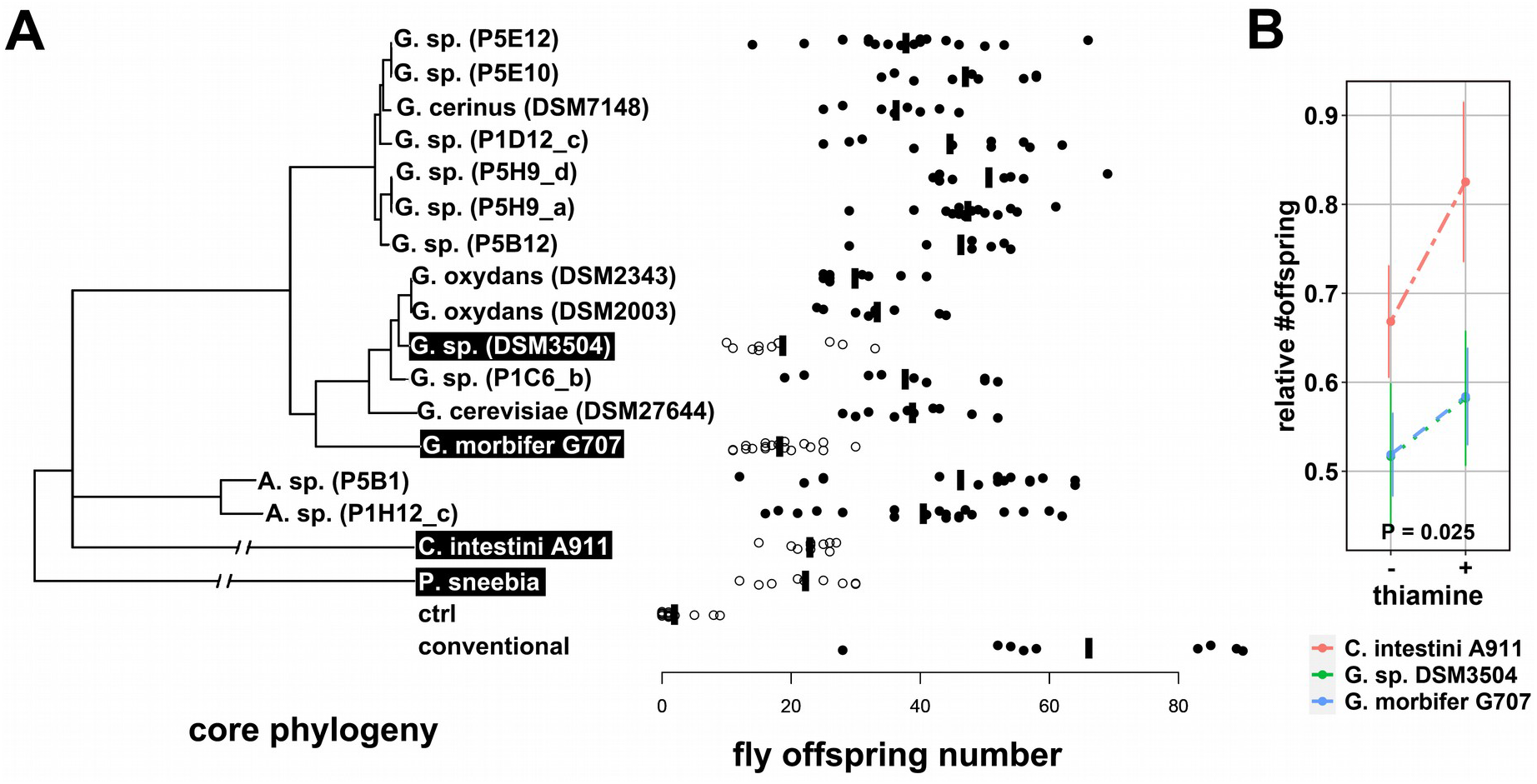
(A) left: Bacterial tree based on 134 single-copy orthologs. Leaf labels of bacteria that do not carry a complete thiamine biosynthesis pathway are on black background. Right: Number of offspring produced by mono-associated CantonS females; vertical bars: median; ctrl: axenic flies; conventional: flies reinfected with lab microbiota. (B) The effect of thiamine treatment (x-axis) on the number of offspring in mono-associated females. Under thiamine treatment, the number of offspring increased for the strains that do not possess a complete thiamine biosynthesis pathway relative to strains that possess the complete pathway. P-value determined by linear mixed effects model. Error bars represent the standard error of the mean.

**Table 1.**
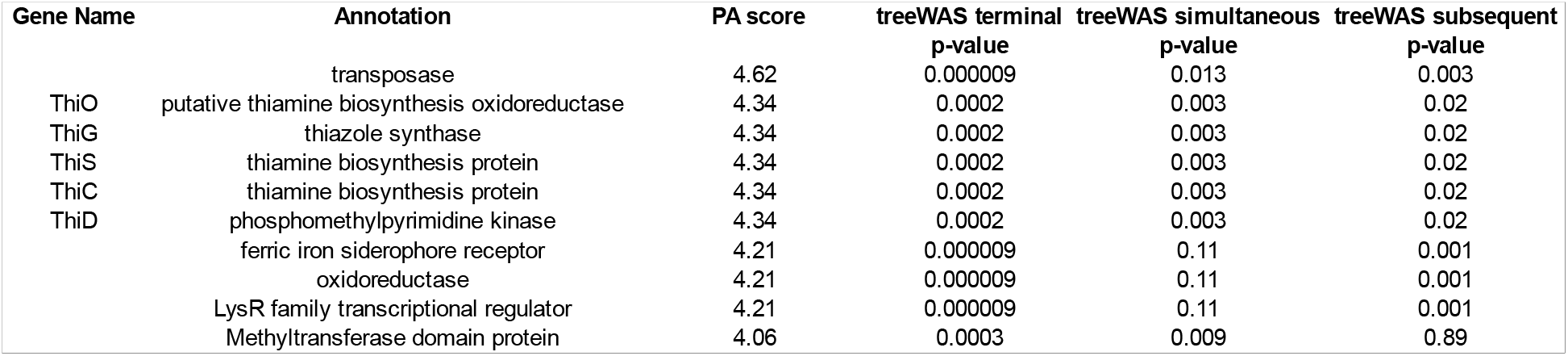
List of the ten genes that were most strongly associated with offspring number according to PA-association scores. All associations were confirmed by at least one of three methods from treeWAS [21].

| Gene Name | Annotation | PA score | treeWAS terminal<br>p-value | treeWAS simultaneous<br>p-value | treeWAS subsequent<br>p-value |
| --- | --- | --- | --- | --- | --- |
|  | transposase | 4.62 | 0.000009 | 0.013 | 0.003 |
| ThiO | putative thiamine biosynthesis oxidoreductase | 4.34 | 0.0002 | 0.003 | 0.02 |
| ThiG | thiazole synthase | 4.34 | 0.0002 | 0.003 | 0.02 |
| ThiS | thiamine biosynthesis protein | 4.34 | 0.0002 | 0.003 | 0.02 |
| ThiC | thiamine biosynthesis protein | 4.34 | 0.0002 | 0.003 | 0.02 |
| ThiD | phosphomethylpyrimidine kinase | 4.34 | 0.0002 | 0.003 | 0.02 |
|  | ferric iron siderophore receptor | 4.21 | 0.000009 | 0.11 | 0.001 |
|  | oxidoreductase | 4.21 | 0.000009 | 0.11 | 0.001 |
|  | LysR family transcriptional regulator | 4.21 | 0.000009 | 0.11 | 0.001 |
|  | Methyltransferase domain protein | 4.06 | 0.0003 | 0.009 | 0.89 |

### The bacterial thiamine biosynthesis pathway is associated with increased offspring number

Five of the six bacterial genes that were most strongly associated with offspring number were part of the thiamine biosynthesis pathway (TBP, Table 1). Females reared on bacteria carrying a complete TBP (TBP+) produced more offspring (*P* = 0.0038, Mann-Whitney-Test on strain medians, n = 17, Figure 1A), suggesting that bacterial thiamine production might increase the number of offspring.

Because high numbers of *Drosophila* offspring on a confined resource like a *Drosophila* vial can lead to crowding effects including smaller adults and reduced individual fitness, we weighed the adult flies at the end of the experiment. Weight did not differ significantly between the offspring of females reared on TBP+ and TBP-strains (*P =* 0.55, Mann-Whitney-Test on strain medians, n = 17, Figure S2), providing no evidence for larval crowding or reduced adult size.

The increase in the number of adult offspring for females reared on TBP+ strains could result from a shortened development time or an increased number of eggs laid by the females. Due to the lack of evidence that bacterial thiamine increases egg laying [37], we expected that time to puparium formation would be shorter for flies associated with TBP+ bacteria. Indeed, pupariation time was significantly reduced for offspring of females reared on TBP+ strains (*P =* 0.015, Mann-Whitney-Test on strain medians, n = 17, Figure S3). Significance of all p-values was confirmed in a linear model framework that accounts for bacterial load and also in a phylogenetic ANOVA (Script S1).

In order to further explore a potential role of thiamine in increasing fly offspring number, we supplemented the diet of females that were mono-associated with seven bacterial strains (*G. oxydans* DSM2343, *G. oxydans* DSM2003, *G. sp.* DSM3504, *G. sp.* P1C6_b, *G. cerevisiae* DSM27644, *G. morbifer* G707 and *C. intestini* A911) with thiamine. This setup covered the *G. morbifer* branch of the phylogeny and all acetic acid bacteria that are missing key enzymes of the TBP (TBP-). The relative number of offspring for the three TBP-strains increased under thiamine treatment (*P* = 0.025, linear mixed effects model, Figure 1B), supporting a role of bacterial thiamine production in the number of offspring that flies produced. Furthermore, with thiamine added, neither average offspring weight nor the time to pupariation differed between TBP+ and TBP-associated flies (*P* = 1 both cases, Mann-Whitney-Test on strain medians, n = 7, Figure S4).

### Thiamine biosynthesis genes were most likely lost and reacquired by horizontal gene transfer as an operon on the branch that includes *G. morbifer*

In order to better understand the evolutionary history of the TBP (Figure 2A) in *Gluconobacter*, we analyzed the synteny of the underlying loci in a phylogenetic framework. The strains in the upper two panels of Figure 2B possess all genes required for thiamine biosynthesis. A closer inspection of TBP genes on the *G. morbifer* branch (II in Figure 2C) revealed that two strains are TBP-, while the four other strains are TBP+. Inspection of the TBP gene loci revealed that all strains on branch II are missing the operon like structure thiOSG (Figure 2C) at the locus that is syntenic with branch I. The same pattern was found for thiC and thiD (Figure S5). thiOSG (Figure 2C), thiC, and thiD (Figure S6) are present in the closely related bacteria *Gluconobacter samuiensis* and *Neokomagateaa tanensis* at syntenic loci, suggesting deletion on branch II. The strains with an intact operon on branch II carried a TBP operon at different loci (Figure 2D), suggesting insertion.

**Figure 2.**
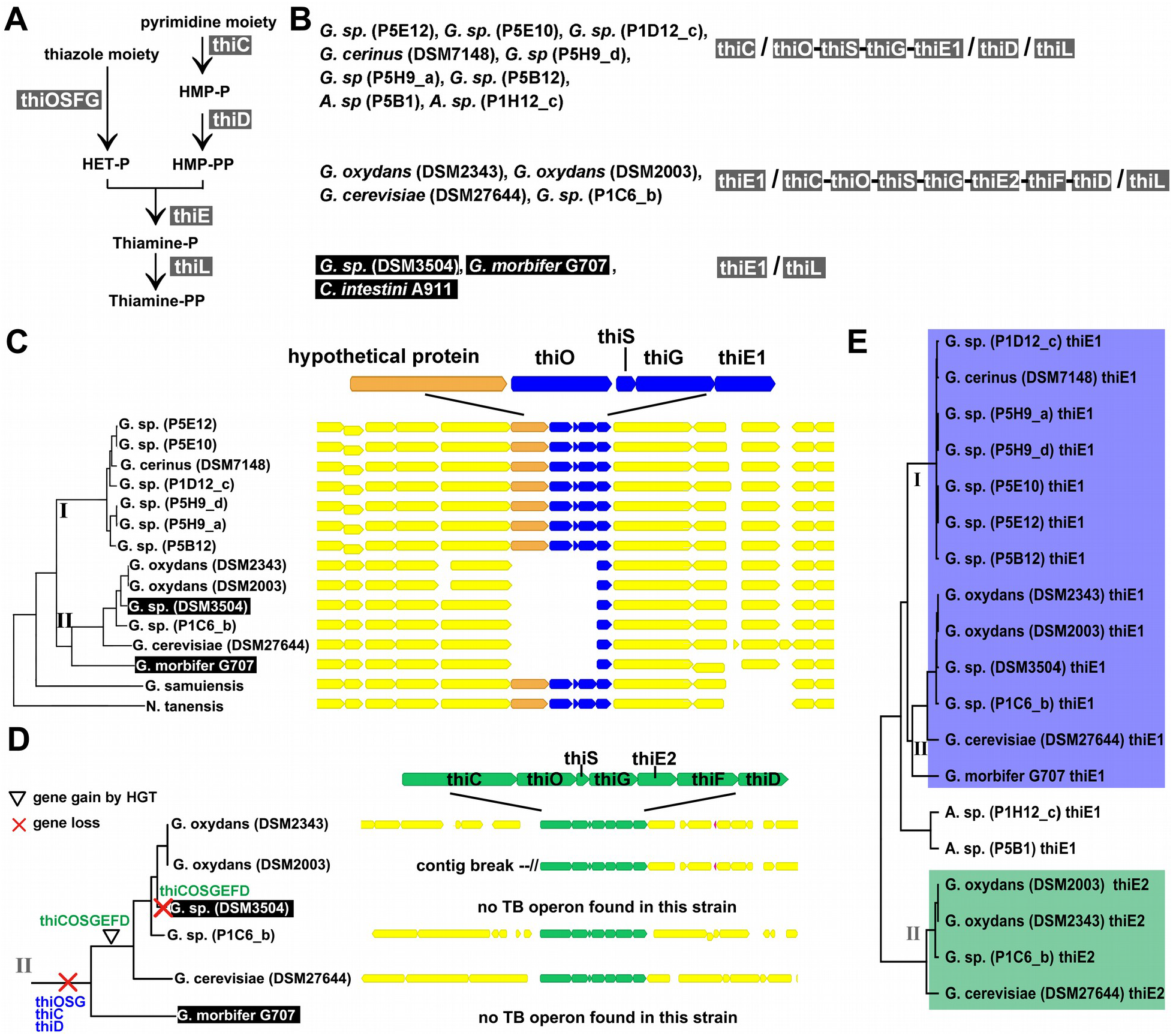
(A) The thiamine biosynthesis pathway in acetic acid bacteria (B) Overview of thiamine biosynthesis genes in the analyzed bacteria. Note that the function of thiF that appears to be missing in the strains of the upper row can be replaced by the function of the homolog MoeB (Rodionov et al., 2002) that we found in all strains analyzed. Genes forming an operon are separated by a hyphen. Genes from different loci are separated by slashes. (C) Synteny of the flanking regions of thiamine biosynthesis genes in *Gluconobacter.* thiOSG are missing on the *G. morbifer* branch (II) at this locus. Thiamine biosynthesis genes are in blue. (D) right: The complete pathway to synthesize Thiamine-P (green) forms an operon on the *G. morbifer* branch (branch II); left: The phylogeny depicts the inferred evolutionary scenario on branch II. (E) Phylogeny of thiE. *G. oxydans* DSM2343, *G. oxydans* DSM2003, *G. sp.* P1C6_b, *G. cerevisiae* DSM27644 have two copies of thiE, thiE1 (blue) and thiE2 (green). The phylogeny of thiE1 (blue background) is congruent with the core genome phylogeny. ThiE2 (green background) forms a distinct clade that is more distant than thiE from *Acetobacter*, indicating HGT from a distant clade.

Analyzing the sequences of the inserted genes in a phylogenetic framework, we found that the inserted genes form a distant clade. For example thiE1, the copy that remained at the locus shown in Figure 2C, followed the phylogeny based on the core genome, while the potentially newly acquired copy thiE2 that is part of the operon thiCOSGEFD formed a distant clade (Figure 2E), supporting HGT. Within this clade, the phylogeny of thiE2 is again congruent with the core genome phylogeny, consistent with a single reacquisition event of thiCOSGEFD. The same phylogenetic patterns were found for the other TBP genes that were shared across branches (thiCOSGD, Figure S7), further supporting a single HGT of thiCOSGEFD to the *G. morbifer* branch. Because TBP genes can occur on plasmids [74], we blast searched the plasmids of the strains for which the plasmids were resolved for TBP genes, finding no evidence for TBP genes (data not shown). In order to identify a potential donor of the operon, we blast searched the sequence of the entire operon against the ncbi non-redundant nucleotide database (nr). The best matching non-*Gluconobacter* sequences were from Rhodobacteraceae, a phylogenetically distant bacterial family (Table S3). A single reacquisition event of the essential TBP genes, as suggested by the concordance of the inserted operon with the core gene phylogeny, implies that the TBP-strain DSM3504 lost the operon again in an independent event, as depicted in Figure 2D (left).

## Discussion

### Microbial GWAS for host traits can benefit from strain level variation

We applied a microbial GWAS approach that associates bacterial genes with host phenotype focusing on the genus *Gluconobacter.* Microbial GWAS approaches can be particularly powerful, when pan-genomic variation of closely related bacterial strains can be leveraged, as has been shown for e.g. virulence genes [75]. We showed that genetic variation between the strains P1C6b, DSM2003, DSM2343, and DSM3504 (mean ANI = 95.5%) empowered us to pinpoint the TBP (Figure S1). Variation between bacteria that have ANI > 95% is considered strain level variation [76]. The only gene that had a higher association score for offspring number than the TBP genes was a transposase. Transposases more frequently produce rare presence-absence patterns because they are mobile and not linked as strongly to the rest of the genome as are non-mobile genetic elements. Therefore, we suspect that the high association score is an artefact of its mobility although we can not exclude an effect of the transposon on the number of fly offspring.

### Variation between closely related microbes is important for host phenotypes

We observed significant variation of phenotypes between flies that were associated with closely related microbial strains. This supports the notion that strain level variation is important to consider when studying host-microbe interaction in animals, humans, and plants alike (e.g. [15, 18, 52, 77, 78]. In particular, in *D. melanogaster* evidence for the importance of variation between closely related bacteria is accumulating for life history of the host [26, 29, 47–51, 79]. Unawareness of strain level variation in bacterial effects on the host might have led to perceived inconsistency between studies [53].

### Loss of the TBP and regain by HGT in the context of the evolution of host-microbe interaction

Fly fitness was strongly associated with genes from the TBP. In [37], thiamine produced by *Acetobacter pomorum* is sufficient to rescue the development of *D. melanogaster* on a thiamine free diet. Furthermore, the authors found that thiamine affects development, but not egg laying nor longevity. This is consistent with our result that flies raised on TBP-strains took longer to develop into pupae (Figure S3). The acquisition of B vitamins like thiamine (B1) is a typical benefit that insects receive from microbes [25, 80] and falls into the greater context of nutrient provisioning by microbes, which is a recurring theme in the evolution of host-microbe interaction [40, 41].

By tracing the genes of the TBP across genomes and the phylogeny, we found that the pathway to produce Thiamine-P was regained most likely via HGT (Figure 2D). As such, our study exemplifies that individual events of HGT into a host-associated microbe can alter host fitness outcomes. Other studies that show an effect on host fitness via HGT to a host-associated microbe involve defensive compounds produced by microbial symbionts in plants [81] and animals [17, 82]. In our study, the increase in host fitness with the reacquisition of the TBP is most likely mediated via nutritional benefits. Only a few similar cases have been described so far. The most prominent may be the acquisition of vitamin B7 (biotin) synthesis by bed bug-associated *Wolbachia* [83]. Similarly, cat flea-associated *Wolbachia* seem to have gained the ability to produce biotin via HGT from closely related strains [84]. In ticks, *pabA* (and possibly *pabB)* required for the synthesis of folic acid, was acquired by a *Coxiella* like symbiont through horizontal gene transfer (HGT) from an Alphaproteobacterium [85] and is thought to affect tick fitness.

A difference between our- and these previous studies is that although *Gluconobacter* is frequently associated with *D. melanogaster* under natural conditions [54–57], it is neither an obligate symbiont nor is it restricted to the fly gut. Given occurence in the environment and the opportunity for horizontal transfer of the bacterium between hosts, there is currently no evidence that the fly host significantly affects *Gluconobacter* evolution. Further, taking into account evidence that the abundance of mobile metabolic genes is governed by selection [86, 87], we must assume that loss and gain of the TBP must first of all benefit the bacterium to persist in the bacterial genome. Thiamine is considered essential for bacteria [88], and thus the TBP can only be lost if enough thiamine is available in the environment. Fruit, the main food substrate of *Drosophila* under natural conditions [89] and the basis for the food used in our study, is mostly poor in thiamine [90]. However, other bacteria that are associated with *Drosophila*, for example other strains of *Gluconobacter* (this study), *Acetobacter pomorum* or *Lactobacillus plantarum*, can produce thiamine [37, 39]. Under these conditions, it might be beneficial for a community member to lose TBP as a result of selection for reduced metabolic expenditure [91]. This is consistent with TBP dependent fitness effects on the host being a byproduct of selection on thiamine production in the microbe.

Our study suggests that HGT to host associated-microbes could quickly increase host fitness. An increase in microbe mediated host fitness should also increase selection pressure on the host to favor that particular microbe that provides an increased benefit [41, 92, 93]. Waterworth et al. [94] suggested that the acquisition of genes to produce a defensive compound via HGT was key to the domestication of a bacterial defensive symbiont in beetles. We speculate that similar scenarios might be plausible for nutritional benefits in *Drosophila* because (i) mechanisms of host selection work efficiently for environmentally acquired bacteria [95–98], (ii) stable, strain specific associations of *Drosophila* with mutualistic bacteria have been reported [50], and (iii) evidence for host selection in the fly is accumulating in the laboratory [45, 99] as well as under natural conditions [55, 57]. Because the result of HGT here provides a potential benefit to the host under thiamine poor conditions that are often encountered under natural conditions e.g. on thiamine poor fruit, our study contributes to a broader view of adaptation that can involve a flexible microbiome [4, 100].

## Supporting information

supplementary figures tables and scripts

## Data availability

Raw sequence data for the bacterial genomes, the assemblies used in this study, and the 16S rRNA gene sequences will be available under PRJNA656529

## Acknowledgements

We thank Ruth Hershberg (Technion, Haifa, Israel), Christian Voolstra (Uni Konstanz, Germany), Lena Waidele (Uni Freiburg, Freiburg, Germany), and John Baines (MPI for Evolutionary Biology, Ploen, Germany) for helpful comments on the manuscript. This work was funded by the DFG (STA1154/4-1; Projektnummer 408908608 and BA5529/1-1; Projektnummer 405974812). The authors acknowledge support by the state of Baden-Württemberg through bwHPC.

## Competing interest

The authors declare no competing interests.

## Notes

### Competing Interest Statement

The authors have declared no competing interest.

