## supplementary figures tables and scripts for "Horizontal gene transfer-mediated bacterial strain variation affects host fitness": ScriptS1stats.R

```
#load package and function
library(MASS)
library(multcomp)
library(geiger)
library(phytools)
library(lme4)
library(ggplot2)
library(Rmisc)
overdisp_fun <- function(model) {
  rdf <- df.residual(model)
  rp <- residuals(model,type="pearson")
  Pearson.chisq <- sum(rp^2)
  prat <- Pearson.chisq/rdf
  pval <- pchisq(Pearson.chisq, df=rdf, lower.tail=FALSE)
  c(chisq=Pearson.chisq,ratio=prat,rdf=rdf,p=pval)
} #test for overdispersion in modelling

#######*********fly offspring********############
#####load fitness assay data  ###fly offspring
fitness_all=read.csv(file="Table S4.csv", header = TRUE, sep = ",")
fitness=fitness_all[fitness_all$thiamine==0,]#pick fitness data
fitness$bacteria=factor(fitness$bacteria)

######Is there a significant difference in fly offspring among all strains?######
fitness.ba=fitness[!fitness$bacteria%in%c('ctrl','conventional'),] #excluding control treatment
droplevels(fitness.ba$bacteria)->fitness.ba$bacteria

#kruskak wallis test 
kruskal.test(offspring~bacteria,fitness.ba)#P=4.193e-15

######Is there a significant difference in fly offspring between TBP+ and TBP- group?######
#add factor for TBP gene presence, 0 or 1
nrow(fitness.ba)#195
nrow(fitness.ba[fitness.ba$bacteria%in% c('G.morbifer','C.intestini','DSM3504','P.Sneebia'),]) #47
fitness.ba$thigene=rep(1,195)
fitness.ba[fitness.ba$bacteria%in% c('G.morbifer','C.intestini','DSM3504','P.Sneebia'),]$thigene=rep(0,47)

##test 1 using median #offspring for each isolate
med=tapply(fitness.ba$offspring,fitness.ba$bacteria,median)
wilcox.test(med[c(1,5,7,8)],med[-c(1,5,7,8)]) #p-value = 0.003777
#there is a significant difference in fly offspring between TBP+ and TBP-

##test 1.2 additional accounting for CFU
#preparing dataframe
med.dat=as.data.frame(med)
med.dat$thigene=as.factor(c(0,1,1,1,0,1,0,0,1,1,1,1,1,1,1,1,1))
cfu.m2=tapply(fitness.ba$loads, fitness.ba$bacteria, function(x) median(x, na.rm=TRUE))
med.dat$CFU=as.data.frame(cfu.m2)
#modelling
med.lm=lm(med~thigene+cfu.m2,med.dat)
summary(med.lm)    #P=0.000139 for thigene
shapiro.test(med.lm$residuals) #residuals are normally distributed
med.lm1=lm(med~cfu.m2,med.dat)
anova(med.lm,med.lm1) #P=0.000139

#test 1.3 additional accounting for the phylogeny with phylogenetic ANOVA
tree=read.newick('strain_tree2.nwk') #tree from panx pipeline http://panx.dscloud.me:8000/Gluconobacter
#distance in tree cannot be 0
#replace 0 with 0.0000000000000000000001 in newick file 
thigene.factor=med.dat$thigene
names(thigene.factor)=names(med)
aov.phylo(med~thigene.factor, tree, nsim=1000) #2e-04
shapiro.test(resid(lm(med~thigene.factor))) #residuals are normally distributed

#test 1.4 accounting for both CFUs and the phylogeny
#run aov.phylo on the residuals accounting for CFUs
x=resid(med.lm1)
aov.phylo(x~thigene.factor, tree, nsim=1000) #0.0013

##conclusion:
##there is a significant difference in fly offspring between TBP+ and TBP- group
##even when we accounting for CFUs and phylogeny

#######combining data in the second experiment where thiamine is supplemented##########
#######and test if the relative fitness of TBP- strains increase on thiamine supplemented food
#load data
fitness_all2<-fitness_all
fitness_all2$bacteria=factor(fitness_all2$bacteria)
fitness_all2$offspring=as.numeric(as.character(fitness_all2$offspring))
fitness_all2=fitness_all2[!fitness_all2$bacteria%in% c('ctrl','conventional'),] #remove ctrl
droplevels(fitness_all2$bacteria)->fitness_all2$bacteria

#add factor for TBP gene presence, 1 precent, 0 absent
nrow(fitness_all2) #259
nrow(fitness_all2[fitness_all2$bacteria%in% c('G.morbifer','C.intestini','DSM3504'),]) #64
fitness_all2$thigene=rep(1,259)
fitness_all2[fitness_all2$bacteria%in% c('G.morbifer','C.intestini','DSM3504'),]$thigene=rep(0,64)

#pick isolates on G.morbifer branch (Branch II)
#these strains were used in both experiments
branch2=fitness_all2[fitness_all2$bacteria%in% c('G.morbifer','C.intestini','DSM3504','P1C6_b','DSM2343','DSM2003','DSM27644'),]
branch2$thigene=factor(branch2$thigene)
branch2$thiamine=factor(branch2$thiamine)
droplevels(branch2$bacteria)->branch2$bacteria

#####calculating relative fitness
#calculating relfit = offspring_TBP-_strains / mean(all TBP+_strains)
#separtely for thiamine0 and thiamine1 cases
TBP.fit<- summarySE(branch2, measurevar="offspring", groupvars=c("thigene","thiamine"))
TBP0=branch2[branch2$bacteria%in%c('DSM3504','G.morbifer','C.intestini'),] #pick TBP- strains
TBP0$bacteria<-droplevels(TBP0$bacteria)
TBP0.fit<- summarySE(TBP0, measurevar="offspring", groupvars=c("bacteria","thiamine"))
TBP0$relfit=TBP0$offspring 
TBP.fit[3,4] #mean of TBP+ strains in initial experiment
TBP.fit[4,4] #mean of TBP+ strains in the experiment that extra thiamine was supplemented
TBP0[TBP0$thiamine==0,]$relfit=TBP0[TBP0$thiamine==0,]$relfit/TBP.fit[3,4]
TBP0[TBP0$thiamine==1,]$relfit=TBP0[TBP0$thiamine==1,]$relfit/TBP.fit[4,4] 

#Model for Figure 1B model testing, treating bacterial isolate as random effect to account for pseudoreplication
lmer=lmer(relfit~thiamine+(1|bacteria),TBP0) 
lmer1=lmer(relfit~1+(1|bacteria),TBP0)
anova(lmer,lmer1) #0.02545
hist(residuals(lmer)) #residuals look good
shapiro.test(residuals(lmer)) #residuals are normally distributed
#conclusion:
#the relative of fitness increases significantly when thiamine is supplemented

#calculating average relative fitness for Figure 1B
TBP0.fit$average_offspring_TBP_positive=c(rep(c(TBP.fit[3,4],TBP.fit[4,4]),3))
TBP0.fit$rel_fit=TBP0.fit$offspring/TBP0.fit$average_offspring_TBP_positive

#calculating sd by error Propagation for Figure 1B error bars
#www.geol.lsu.edu/jlorenzo/geophysics/uncertainties/Uncertaintiespart2.html
#calculating standard deviation using equation dz/z=((dx/x)^2+(dy/y)^2)^(1/2)
TBP0.fit$average_offspring_TBP_positive_sd=c(rep(c(TBP.fit[3,5],TBP.fit[4,5]),3))
TBP0.fit$rel_fit_sd=((TBP0.fit$sd/TBP0.fit$offspring)^2+(TBP0.fit$average_offspring_TBP_positive_sd/TBP0.fit$average_offspring_TBP_positive)^2)^(1/2)*TBP0.fit$rel_fit
TBP0.fit$rel_fit_se=TBP0.fit$rel_fit_sd/(TBP0.fit$N^(1/2))
#making plot
p<- ggplot(TBP0.fit,aes(x=thiamine, y=rel_fit,group=bacteria,color=bacteria
                     # ,ymin=0, ymax=1
))+
  geom_line(aes(linetype=bacteria),alpha=0.9,size=1.1) +
  geom_point(size=2)+
  scale_linetype_manual(values=c("twodash", "dotted","dashed"))+
  geom_errorbar( aes(ymin=rel_fit-rel_fit_se, ymax=rel_fit+rel_fit_se),width=.02,  position=position_dodge(0.08))+
  labs(title="",y='relative #offspring')+
  theme(plot.title = element_text(size=17.5,face="bold",colour = "black",hjust = 0.5),
        axis.text=element_text(size=16,face="bold",colour = "black"),
        axis.title=element_text(size=17.5,face="bold"),
        legend.text = element_text(size=15,face="bold",colour = "black"),
        panel.border = element_rect(colour = "black", fill=NA, size=1),
        panel.background = element_rect(fill = 'white', colour = 'red'),
        panel.grid.major =element_line(colour = 'grey'))+
  scale_x_discrete(labels = c("-", "+")) +
  scale_colour_discrete(name="",labels = c("C. intestini A911","G. sp. DSM3504", "G. morbifer G707"))+
  guides(linetype=FALSE)+
  annotate(geom="text", x=1.5, y=0.45, label="\nP = 0.025",
           color="black",size=5,fontface="bold")
print(p)


#######*********fly weight and pupariation time********############
#prepare data
#calculating pupation time 
for(i in 1:nrow(fitness_all2)) {
  fitness_all2$time[i]=(fitness_all2[i,6]*6+fitness_all2[i,7]*7+fitness_all2[i,8]*8+fitness_all2[i,9]*9+fitness_all2[i,10]*10+fitness_all2[i,11]*11+fitness_all2[i,12]*12+fitness_all2[i,13]*13+fitness_all2[i,14]*14+fitness_all2[i,15]*15)/sum(fitness_all2[i,6:15])
  fitness_all2$weights[i]=sum(fitness_all2[i,6:15])
}
fitness2=fitness_all2[fitness_all2$thiamine==0,]#pick data in initial experiment

##Is there a significant difference in pupariation time  between TBP+ and TBP-?
#test 1 using median
med=tapply(fitness2$time,fitness2$bacteria,median)
wilcox.test(med[c(1,5,7,8)],med[-c(1,5,7,8)])#0.01513

#test 1.2 using median controlling for CFUs
med.dat=as.data.frame(med)
med.dat$thigene=as.factor(c(0,1,1,1,0,1,0,0,1,1,1,1,1,1,1,1,1))
med.dat=med.dat[order(rownames(med.dat)),]
med.dat$CFU=as.vector(cfu.m2)
#modelling
med.lm=lm(med~thigene+cfu.m2,med.dat)
summary(med.lm)      #P =0.016 for thigene
shapiro.test(med.lm$residuals) #residuals look good
med.lm1=lm(med~cfu.m2,med.dat)
anova(med.lm,med.lm1)     #0.01597

#test 1.3 account for phylogeny, using aov.phylo
thigene.factor=med.dat$thigene
names(thigene.factor)=names(med)
aov.phylo(med~thigene.factor, tree, nsim=1000) #0.0459 around the 0.05
shapiro.test(resid(lm(med~thigene.factor))) #residuals look good

#conclusion
#there is a significant difference in pupariation time  between TBP+ and TBP- strains
#holds true if we control for CFUs or the phylogeny 

###fly weight###
fitness2$weight_per_fly=as.numeric(as.character(fitness2$weight_per_fly))

##Is there a significant difference in fly weight  between TBP+ and TBP-?
#test using median
med=tapply(fitness2$weight_per_fly,fitness2$bacteria,median)
wilcox.test(med[c(1,5,7,8)],med[-c(1,5,7,8)])#0.5487
#no, weight  is not significantly different between TBP+ and TBP- strains in initial experiment

######fly weight and pupariation time## when supplementing thiamine####
#picking data
resc=fitness_all2[fitness_all2$thiamine==1,]
resc$bacteria<-droplevels(resc$bacteria)

##Is there a significant difference in fly pupariation time between TBP+ and TBP- when thiamine was added?
# test using medium
med=tapply(resc$time,resc$bacteria,median)
wilcox.test(med[c(1,5,6)],med[-c(1,5,6)]) #p-value = 1

##Is there a significant difference in fly weight between TBP+ and TBP- when thiamine was added?
resc2=resc[!is.na(resc$weight_per_fly),]#remove one sample without weight data
resc2$weight_per_fly=as.numeric(as.character(resc2$weight_per_fly))
# test using medium
med=tapply(resc2$weight_per_fly,resc2$bacteria,function(x) median(x, na.rm=TRUE))
wilcox.test(med[c(1,5,6)],med[-c(1,5,6)]) #p-value = 1

#conclusion:
#with thiamine added, neither average offspring weight nor the time to pupariation differed between TBP+ and TBP- associated flies
#while in the initial experiment, the time to pupariation differed significantly between TBP+ and TBP- associated flies
```
