## supplementary figures tables and scripts for "Horizontal gene transfer-mediated bacterial strain variation affects host fitness": Supplement.pdf

Overview:

Figure S1 distribution of PA scores

Figure S2 offspring weight

Figure S3 pupariation time

Figure S4A pupariation time with thiamine added to the diet

Figure S4B offspring weight with thiamine added to the diet

Figure S5 thiC and thiD missing at syntenic loci

Figure S6 synteny in *Swingsia samuiensis* and *Neokomagateaa tanensis* for thiC, thiD, and thiOSG

Figure S7 phylogeny of the HGT TBP genes thiC, thiD, and thiOSG

Figure S8 contamination control using 16S rRNA gene sequencing after the experiment

Table S1 list of bacterial strains used in the experiments including assembly information

Table S2 full PA and Treewas results table

Table S3 blast results for HGT operon

Table S4 CFUs (Yun) missing: explanation of columns

Script S1 statistical analyses

Script S2 16S rRNA gene sequence analysis with mothur

Script S3 microbial GWAS

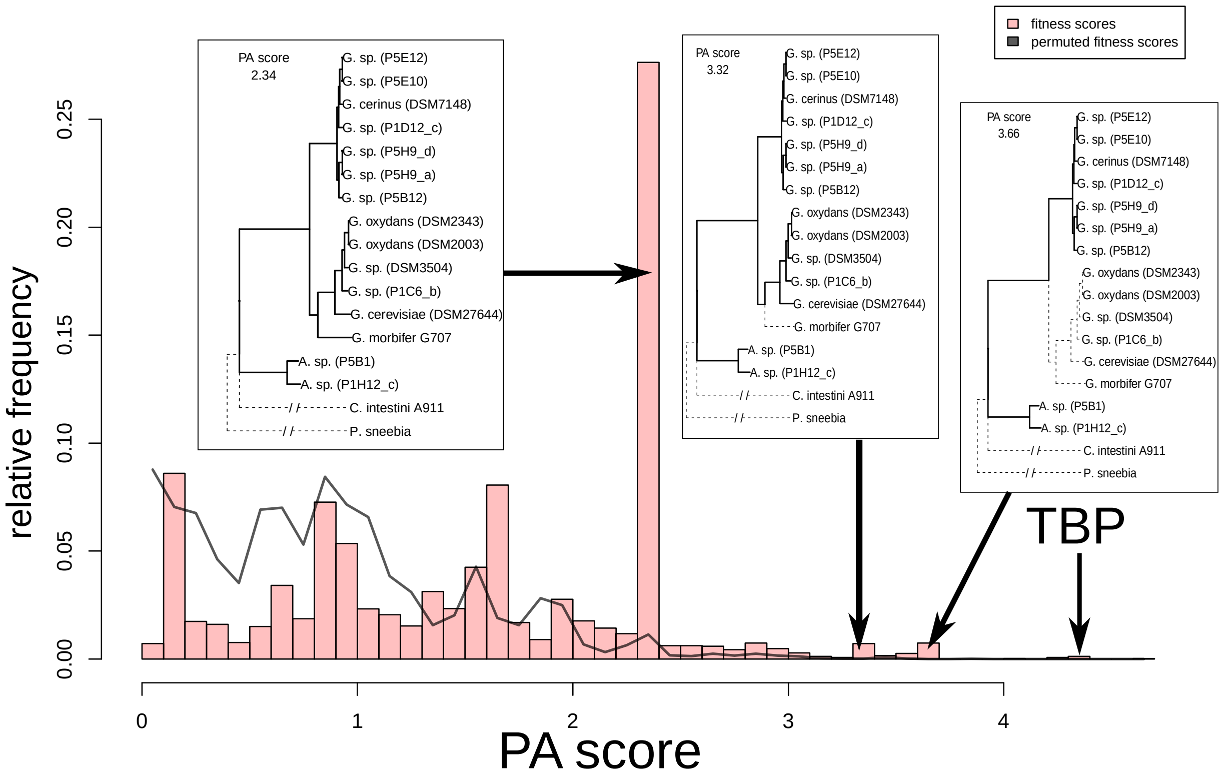


**Figure S1** Distribution of PA-scores. The presence absence score (PA score) for all *Gluconobacter* clusters of homologous genes is shown as a bar chart. The TBP genes have a score of 4.34. The peak at 2.34 corresponds to the genes that are missing in the outgroup strains *C. intestini* and *P. sneebia*, but present in all other strains (phylogeny to the left). The peak at 3.32 corresponds to the genes that are missing in *C. intestini, P. sneebia,* and *G. morbifer* G707. The peak at 3.66 represents the genes that are private to the *G. cerinus* clade and shared with *Acetobacter*. Only with strain variation within the *G. morbifer*/G. *oxydans* clade as shown in Figure 1, we get the extreme PA scores for the TBP. The solid line represents the PA score distribution for 10 datasets with randomly permuted fitness values. We found one cluster with an association score higher than 4.34 in the permuted data sets. This corresponds to the PA scores for the TBP genes being in the 99.999 percentile. The mean PA score for TBP genes from permutations is 0.86 +/- 0.59.

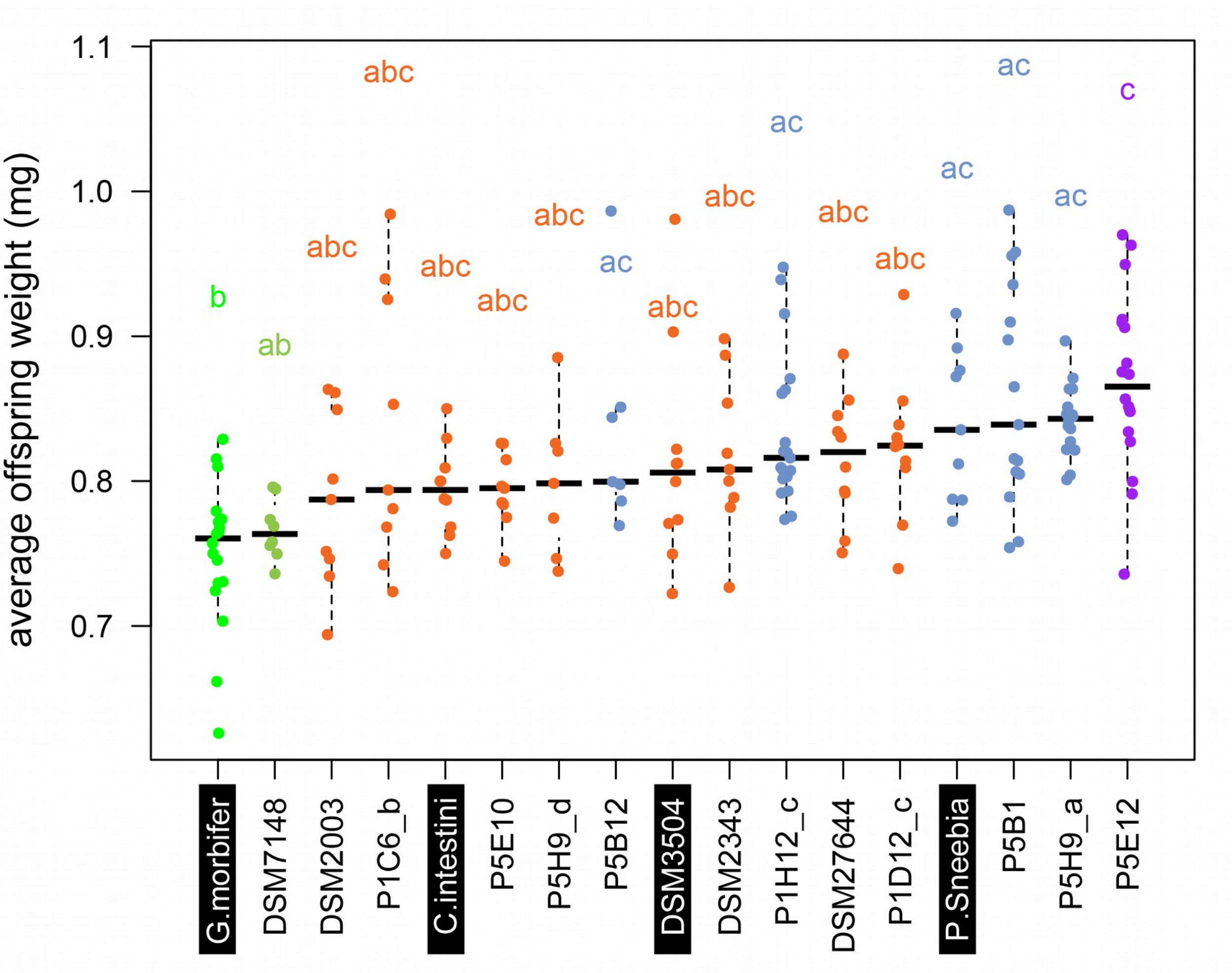


**Figure S2** Weight per offspring from mono-associated female flies. Offspring weight did not differ significantly between TBP+ and TBP- strains. Isolates that do not share one of the letters above (a,b,c) differ significantly according to Tukey's test.

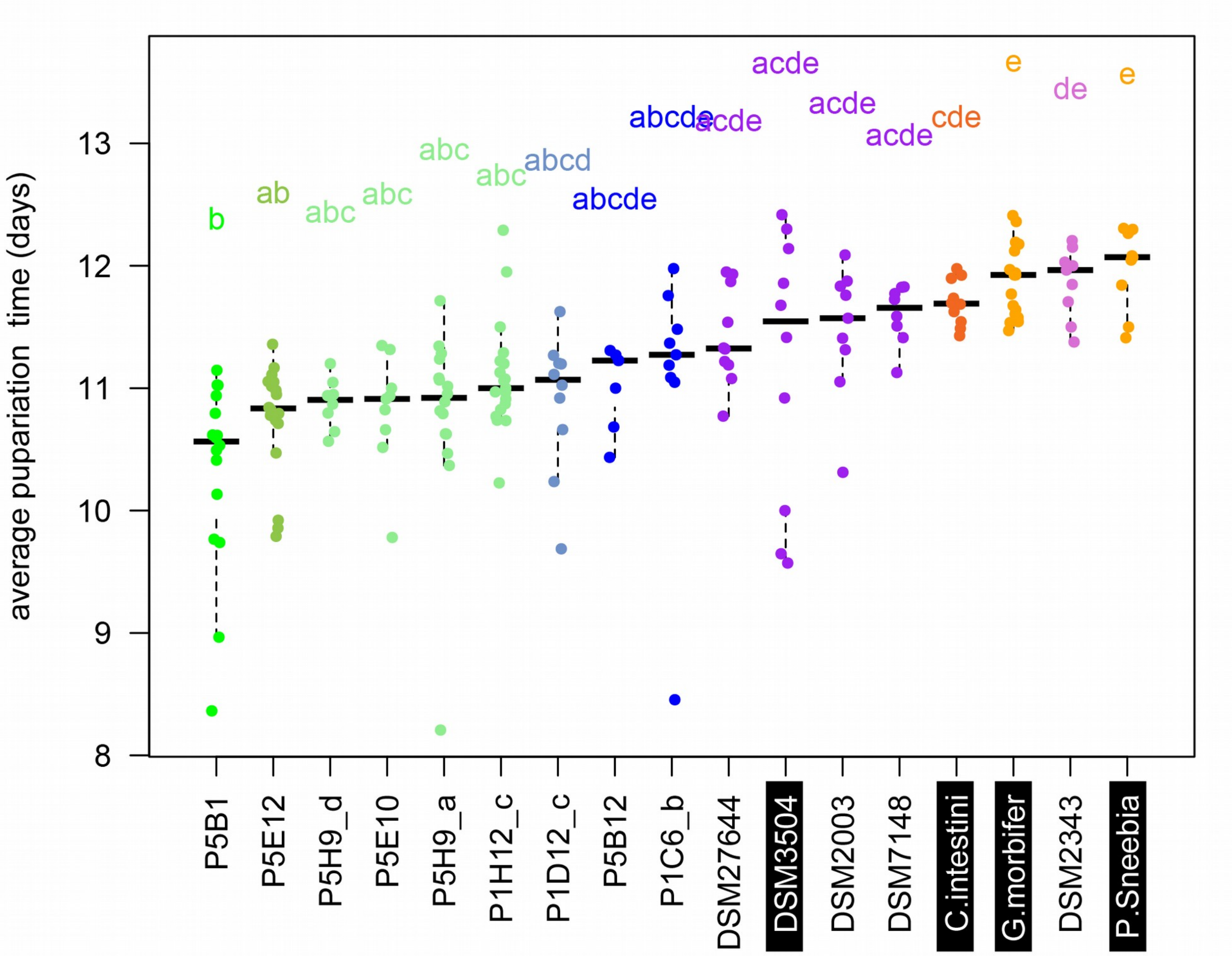


**Figure S3** Average pupariation time of offspring for mono-associated females. Time to pupariation was longer for TBP- strains. Isolates that do not share one of the letters above (a,b,c) differ significantly according to Tukey's test. TBP- strain x-axis labels are on black background.

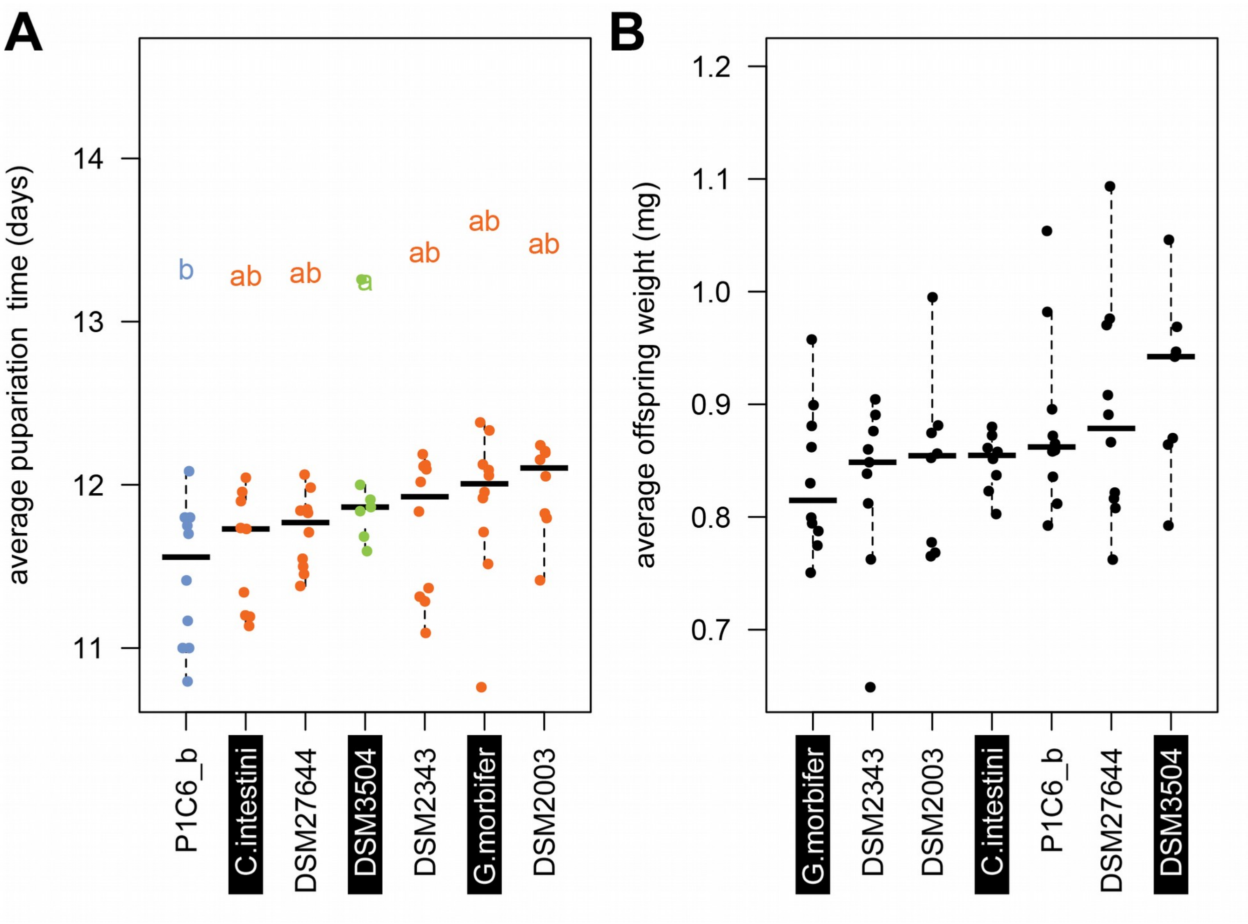


**Figure S4** (A) Average pupariation time of offspring for mono-associated females when the diet was supplemented with 1 µg/ml thiamine. Time to pupariation did not differ significantly between TBP+ and TBP- strains. Isolates that do not share one of the letters above (a,b) differ significantly according to Tukey's test. (B) Weight per offspring from mono-associated female flies when the diet was supplemented with 1 µg/ml thiamine. Offspring weight did not differ significantly between strains.

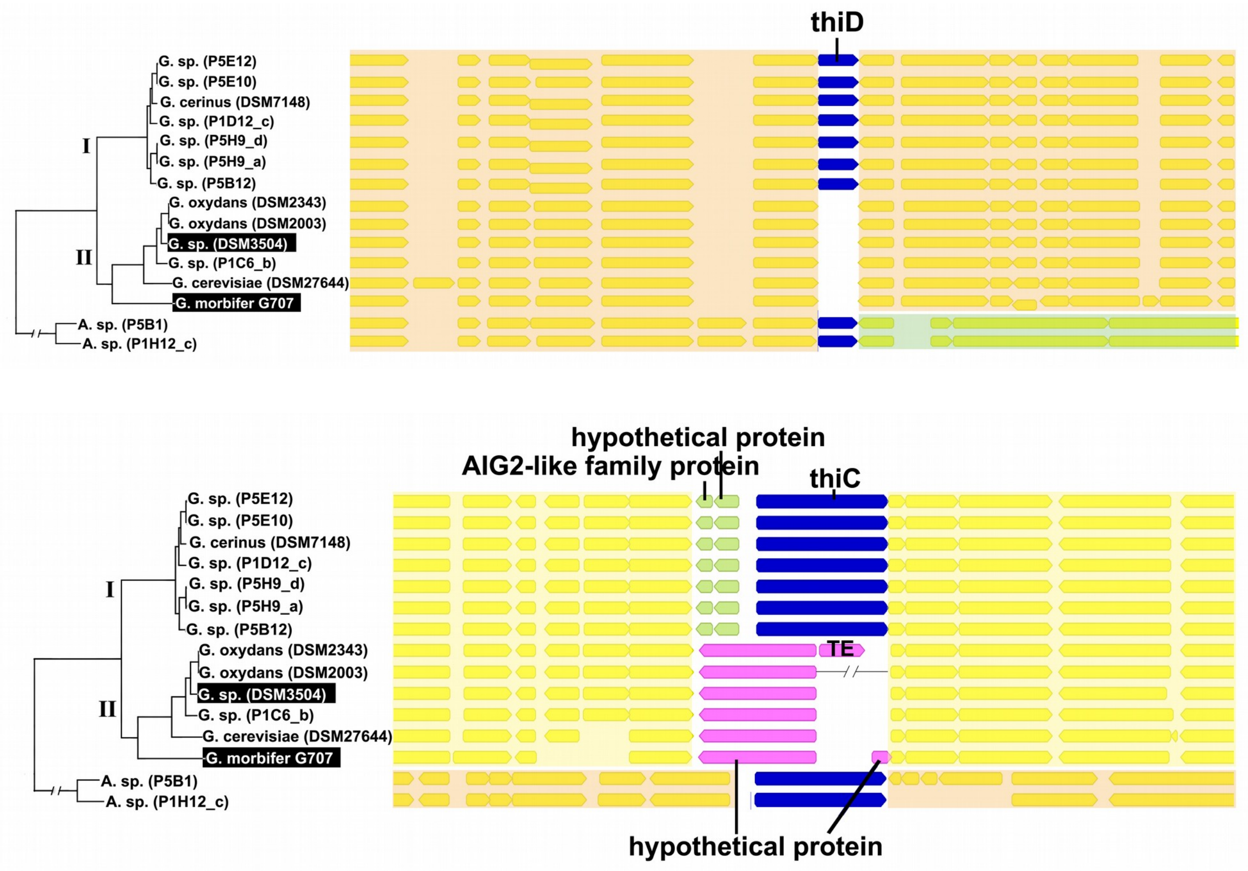


**Figure S5** thiC and thiD are missing at syntenic loci on the *G. morbifer* branch.

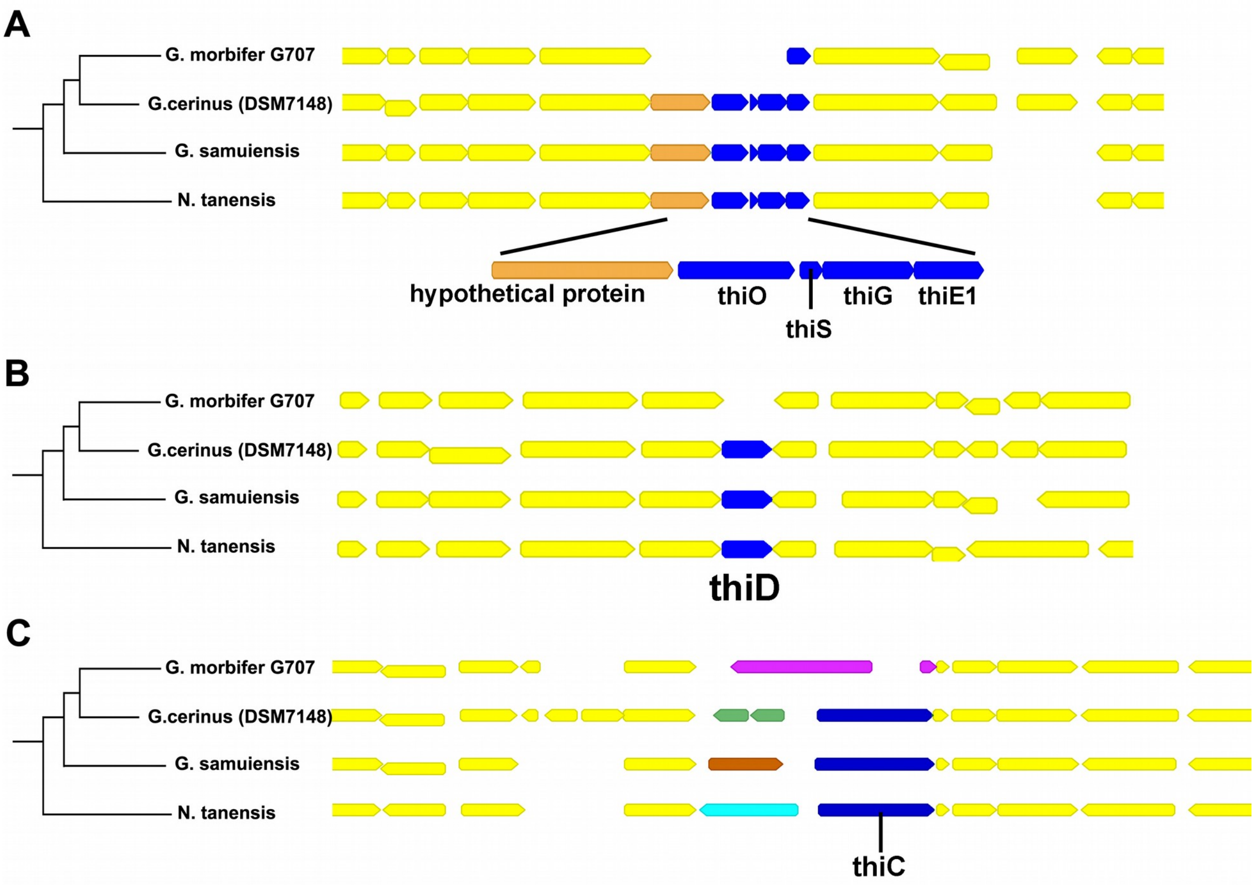


**Figure S6** The closely related species *G. samuiensis* (also called *Swingsia samuiensis*, NZ_CP038141) and *Neokomagateaa tanensis* (NZ_CP032485) possess thiOSG at the syntenic locus pointing towards a deletion on the *G. morbifer* branch. The phylogeny is based on GTDB v95 (Chaumeil et al. 2020). We find the same pattern for thiC and thiD.

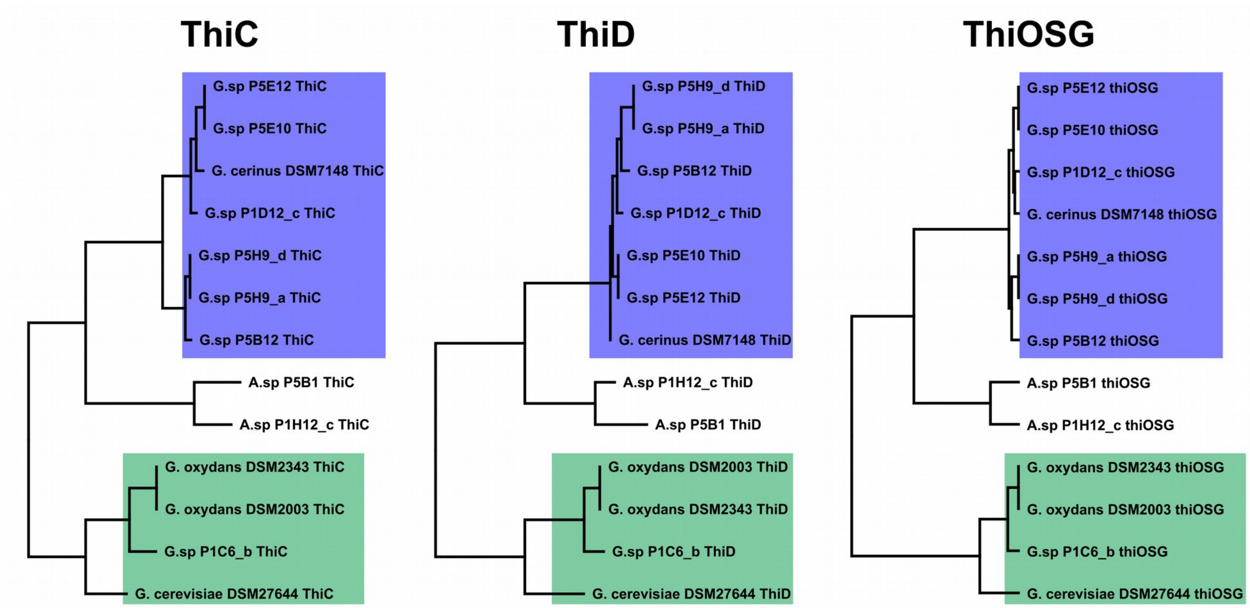


**Figure S7** The phylogeny of the TBP genes (thiC, thiD, thiOSG) is incongruent with the core phylogeny. The TBP genes that were acquired by HGT in the strains G. oxydans DSM2343, G. oxydans DSM2003, G. sp. P1C6_b, and G. cerevisiae DSM27644, form a distant clade (green background). Within this clade the phylogeny is congruent with the core phylogeny further supporting a single HGT of the operon to the *G. morbifer* branch (branch II in Figure 2). The phylogenies of thiC and thiD were extracted from the Panx pipeline (Ding et al., 2018). For the phylogeny of thiOSG, nucleotide sequences were aligned using MUSCLE v3.8.425 (Edgar, 2004) and the tree was built using MrBayes 3.2.6 (Ronquist et al., 2012) as incorporated in Geneious Primer 2020.1.1 (Biomatters ltd.).

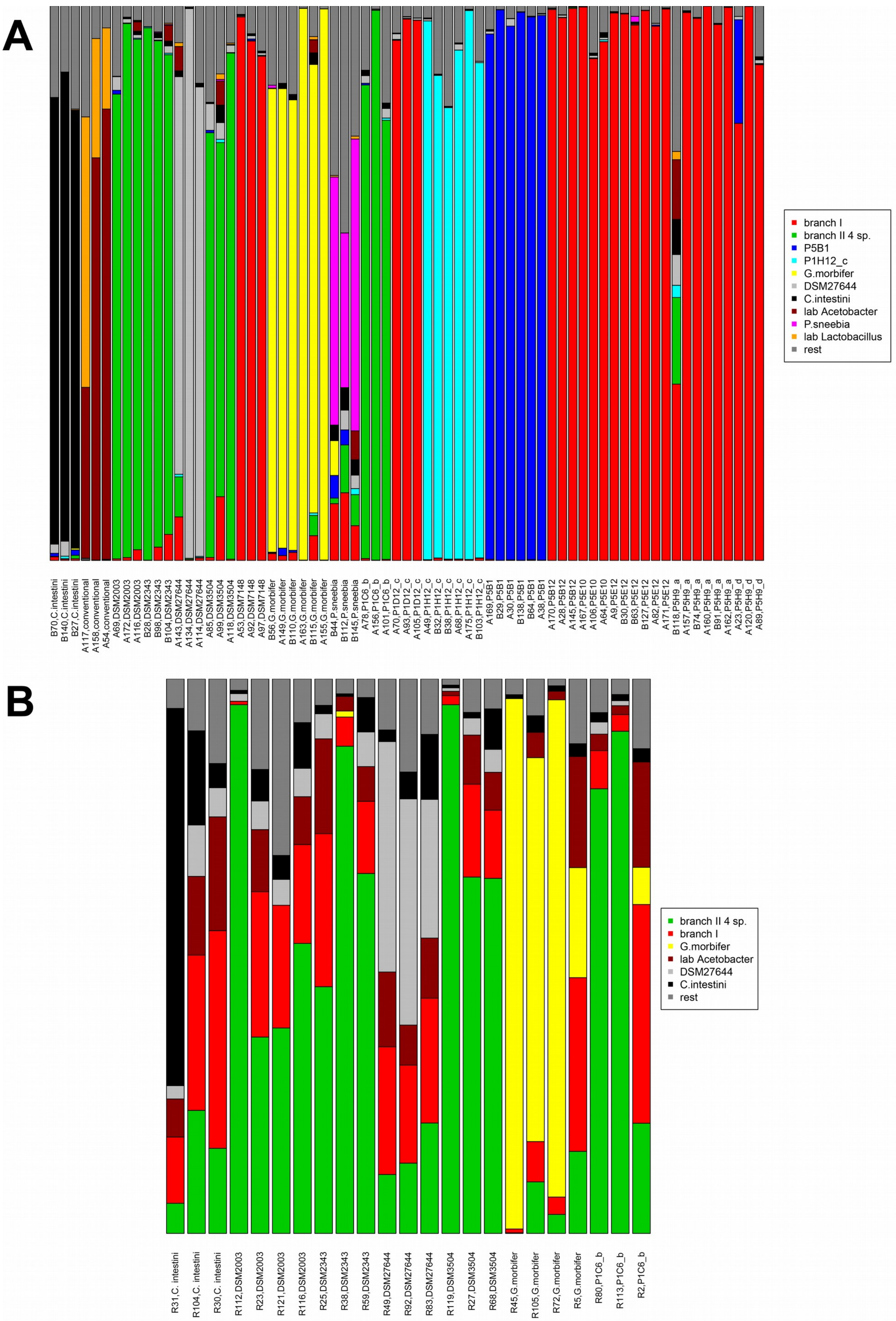
A
branch I
branch II 4 sp.
P5B1
P1H12_c
G.morbifer
DSM27644
C.intestini
lab Acetobacter
P.sneebia
lab Lactobacillus
rest
B70,C.intestini
B140,C.intestini
B27,C.intestini
A117,conventional
A158,conventional
A54,conventional
A69,DSM2003
A172,DSM2003
A116,DSM2003
B28,DSM2343
B98,DSM2343
B104,DSM2343
A143,DSM27644
A134,DSM27644
A114,DSM27644
A85,DSM3504
A99,DSM3504
A118,DSM3504
A53,DSM7148
A92,DSM7148
A97,DSM7148
B56,G.morbifer
A149,G.morbifer
B110,G.morbifer
A163,G.morbifer
B115,G.morbifer
A155,G.morbifer
B44,P.sneebia
B112,P.sneebia
B145,P.sneebia
A78,P1C6_b
A156,P1C6_b
A101,P1C6_b
A70,P1D12_c
A93,P1D12_c
A105,P1D12_c
A49,P1H12_c
B32,P1H12_c
B38,P1H12_c
A68,P1H12_c
A175,P1H12_c
B103,P1H12_c
A169,P5B1
B29,P5B1
A30,P5B1
B138,P5B1
B64,P5B1
A38,P5B1
A170,P5B12
A28,P5B12
A145,P5B12
A167,P5E10
A106,P5E10
A64,P5E10
A9,P5E12
B30,P5E12
B63,P5E12
B127,P5E12
A82,P5E12
A171,P5E12
B118,P5H9_a
A157,P5H9_a
B74,P5H9_a
A160,P5H9_a
B91,P5H9_a
A162,P5H9_a
A23,P5H9_d
A120,P5H9_d
A89,P5H9_d
B
branch II 4 sp.
branch I
G.morbifer
lab Acetobacter
DSM27644
C.intestini
rest
R31,C. intestini
R104,C. intestini
R30,C. intestini
R112,DSM2003
R23,DSM2003
R121,DSM2003
R116,DSM2003
R25,DSM2343
R38,DSM2343
R59,DSM2343
R49,DSM27644
R92,DSM27644
R83,DSM27644
R119,DSM3504
R27,DSM3504
R68,DSM3504
R45,G.morbifer
R105,G.morbifer
R72,G.morbifer
R5,G.morbifer
R80,P1C6_b
R113,P1C6_b
R2,P1C6_b

**Figure S8** The relative abundance of bacterial taxa on flies after the experiments. (A) For the initial fitness assay. The average relative abundance of target 16S sequences was over 88% in the fitness assay, indicating successful inoculation with the target bacterium. Only in 6 out of 66 replicates, the relative abundance was below 75%, including 3 cases of *P. sneebia* that showed low bacterial loads. We think that *P. sneebia* did not grow as well as the other strains under our experimental conditions. (B) For the thiamine supplement experiment. The target isolates were significantly enriched in the replicates (P in the range of 0.000082~0.028, Mann-Whitney-Test). However, we found potential contamination (red and brown bars). The two samples (R112 and R45) from an independent PCR and purification round showed significantly less contamination, suggesting that the contamination might have taken place during the PCR or sequencing steps and not during the experiment.

Three to six replicates per bacterial strain were selected for sequencing as described in the main text. Bacterial communities were profiled by 16s rRNA gene sequencing of whole homogenized flies following Kozich et al. 2013 and Wang et al. 2018. A detailed analysis script can be found as script S2 in the supplementary files.
